# Uptake and Implementation of Multiverse-style Analyses Across 613 Studies

**DOI:** 10.64898/2026.07.15.738584

**Authors:** Andrew Nepomuceno, Angikar Ghosal, Alejandro Sandoval-Lentisco, John P. A. Ioannidis

## Abstract

Empirical conclusions can depend on the many individual choices researchers make when analyzing data. Multiverse-style analyses address this by computing results across a set of specifications rather than a single one, but it is unclear how widely they are being used and how they are being implemented. We searched the Web of Science Core Collection (May 2026) for articles citing six foundational papers on different variants of multiverse-style methods and classified each of the citing articles as implementing such methods or only discussing them. For implementations, we recorded the framework used, the number of specifications, which of four decision nodes (measurement, data processing, modeling, and estimation) were varied, and other aspects such as how results were visualized and interpreted. Of the 1545 classifiable articles, 613 (39.7%) implemented a multiverse-style analysis (primarily multiverse n = 336, specification curve n = 175, vibration of effects n = 20, multimodel n = 59, and multi-analyst/many-analyst n = 23). Uptake spanned many disciplines, most often psychology (39.8%), the social sciences (20.7%), and medicine (19.2%). The number of specifications ranged from fewer than ten to more than ten thousand (median = 144, IQR 24–1248). Modeling (75%) and data-processing (62%) choices were included most often. Interpretation was predominantly descriptive, whereas formal inference (10.5%), preregistration (8.6%), and explicit attention to the defensibility of specifications (3.9%) were uncommon. Multiverse-style analyses are increasingly becoming established across the quantitative sciences, but some implementation practices can be strengthened so that these analyses become more fully transparent and more genuinely informative about the robustness of research findings.

---

Empirical findings depend not only on the data but also on the many analytical choices a research team makes: how constructs are defined and measured, how the raw data are processed, which statistical model is fitted, and how its parameters are estimated. Many of these choices are made from several alternatives, which may be defended in various ways by the investigators, yet most published papers report only one of many possible analytical routes. This problem, often described with terms such as “researcher degrees of freedom” or the “garden of forking paths”, can substantially affect the results, because paths can lead to different, and sometimes opposing, conclusions (Gelman & Loken, 2013; Simmons et al., 2011). The extent of analytical options is considered to be one of the most influential factors that diminish the reliability and trustworthiness of the published scientific literature, especially when coupled with selective reporting (Ioannidis, 2005). The space of reasonable analyses can be enormous, and in some data-rich fields they may encompass thousands of distinct pipelines for a single research question.

Over the past two decades, reforms such as data and code-sharing mandates, and preregistration have tried to make research more transparent and its results easier to check (Munafò et al., 2017; Nosek et al., 2018). Most reforms, however, largely target repeatability and replicability. Repeatability (a.k.a. methods reproducibility (Goodman et al., 2016)) asks whether a reported result can be recomputed from the same data and code, and a major stumbling block is the unavailability of these resources (Brodeur et al., 2026; Ioannidis et al., 2009). Replicability (a.k.a. results reproducibility (Goodman et al., 2016)) addresses whether a finding recurs in new data (Tyner et al., 2026). However, a result can be both repeatable and replicable and yet still hinge on one analyst’s particular choices (Goodman et al., 2016; Nosek et al., 2022). Assessing this third concern, i.e., whether a conclusion is robust to the analytical decisions that could have been made otherwise, is a distinct and arguably deeper task. Its importance is underscored, for example, by many-analyst studies, which repeatedly find that independent teams given the task of analyzing the same data and question often adopt different analytical paths and may reach markedly different conclusions (Botvinik-Nezer et al., 2020; Breznau et al., 2022; Silberzahn et al., 2018).

Different variants of multiverse-style analyses were developed as a systematic response to this problem. Rather than treating alternative analyses as informal, ad hoc sensitivity checks, they ask researchers to enumerate a wide set of specifications, compute the result of each, and report the resulting distribution, often displayed graphically and, in some implementations, tested jointly against a null hypothesis (e.g., Simonsohn et al., 2020). Several influential frameworks share this logic: the vibration of effects (Patel et al., 2015), multiverse analysis (Steegen et al., 2016), multimodel analysis (Young & Holsteen, 2017), multi-analyst (many analysts) studies (Aczel et al., 2021; Silberzahn et al., 2018), and specification curve analysis (Simonsohn et al., 2020). These various approaches emerged in different disciplines and initially might have emphasised slightly different parts of the analytical pipeline; e.g., Steegen et al.’s multiverse focused on data-processing decisions, whereas Young and Holsteen’s multimodel analysis focused on alternative model specifications, although the multiverse logic can in principle be applied across any of the decision nodes when conducting a study and its analysis. Many aspects of how these analyses should be conducted, moreover, remain under active development, including how best to justify the set of specifications examined, how to visualize the results, and how to draw valid inferences from them, as a recent review discusses (Short et al., 2026).

Despite the growing attention these methods have received, it remains unclear how often they have actually been adopted, as opposed to merely being cited, and how they are implemented when they are used. A citation may reflect the genuine deployment of a multiverse-style analysis or only a passing mention of the idea within a broader discussion. Equally little is known about the practical characteristics of deployments: how many specifications are examined, which parts of the analytical pipeline are actually varied, whether the results are visualized, and whether the specification set is interpreted descriptively or with formal inference. Because the value of a multiverse may eventually depend on how it is constructed and interpreted, these implementation details may bear directly on what such analyses can support.

In this meta-research study, we aim to map the uptake and several aspects of the implementation of multiverse-style analyses across the literature that cites six foundational papers. We classify each citing article as either deploying a multiverse-style analysis or only discussing it, and we describe the fields in which these methods are applied and the frameworks that predominate. For the studies that deployed such an analysis, we then characterize different aspects of how it was implemented; the number of specifications examined, which of the decision components were varied, how the results were displayed and, in a subsample coded in greater depth, additional features such as preregistration, whether authors discussed the defensibility and equivalence of the different specifications when defining their multiverses, and how the specification set results were ultimately interpreted.

## Methods

### Search strategy and citation set

In May 2026, we searched the Web of Science Core Collection for all papers that cited one of six foundational articles representing the main multiverse-style methods. These are the most highly-cited papers retrieved on a Web of Science search with the specific names of these methodological variants: “multiverse” analysis (Steegen et al., 2016), “specification curve” analysis (Simonsohn et al., 2020), “vibration of effects” (Patel et al., 2015), “multimodel” analysis (Young & Holsteen, 2017), and “multi-analyst” (“many analysts”) studies, for which we used two foundational articles, one with the term “multi-analyst” and one with the term “many analysts” which was an earlier term for the very same concept (Aczel et al., 2021; Silberzahn et al., 2018). We relied on the literature citing these seminal, highly cited articles rather than on a keyword search because we wanted to target work that explicitly engages with these methods; this approach, however, may miss analyses that implement equivalent ideas without citing these works. According to the Web of Science Core Collection, these articles had received 2,161 citations in total as of May 2026. After deduplication, the working citation set contained 1559 papers. From the Web of Science we also retrieved, for each article, its Research Areas and its citation count (based on all Web of Science databases, which include preprint databases). The Python script used to retrieve the citations is available at https://osf.io/j7nsf/.

### Implementation/discussion classification

We classified each of the 1559 citing papers as either implementing a multiverse-style analysis (*deploy*) or only discussing or citing such methods without implementing them (*discuss*); a further 14 papers could not be accessed and were excluded. Three authors (A.N., A.G., and A.S.L.) first piloted the classification rules on a random sample of 90 citing papers. We then evaluated an LLM (Gemini 3.1 Flash-Lite) as an aid to classification, which reached an accuracy of 92% (80 of 87 papers classified in agreement with manual coding). We used this model to classify the remaining papers, with one author (A.S.L.) manually reviewing every case. The Python script and the exact prompt used for the model queries are available at https://osf.io/j7nsf/.

### Implementation of multiverse-style methods

For the studies classified as *deploy*, we extracted a set of features characterising how each multiverse-style analysis was implemented. Extraction was carried out from the full text and, if necessary, the supplementary materials of all the implemented studies (n = 613). For each study we extracted the framework used (multiverse analysis, specification curve analysis, vibration of effects, multimodel analysis, or a multi-analyst study), the number of specifications reported (where an explicit count could be extracted, n = 529), and which categories of analytical decision were varied across specifications. Following the taxonomy of Hoffmann et al. (2021) and Short et al. (2026), we coded four decision nodes; measurement, data processing, modeling, and estimation, as either varied or not varied in each study. We also extracted whether the results were accompanied by at least one visualization and, if so, its type or types.

A randomly selected subsample of 152 studies (one quarter of the studies classified as deploying a multiverse) was coded in greater depth by one author (A.S.L.). For these studies we additionally extracted other key implementation information, including several points highlighted in Short et al. (2026): whether the multiverse was preregistered; whether the authors discussed the defensibility/equivalence of the competing specifications in the sense of Del Giudice & Gangestad (2021); whether the multiverse-style analysis was the study main analysis or played a secondary role; whether the specifications were fully crossed (nested) or only computed sequentially; whether the authors themselves described the analysis as a “limited” or “mini” multiverse; and how the results were interpreted (considering only descriptive analyses, explicit vote counting, permutation-based inference, or other quantitative methods). The full extraction template and the coded data are available at https://osf.io/j7nsf/.

## Results

### Uptake of multiverse-style analyses

As of May 2026, a total of 1559 articles cited at least one of the target multiverse-style methodological articles (Aczel et al., 2021; Patel et al., 2015; Silberzahn et al., 2018; Simonsohn et al., 2020; Steegen et al., 2016; Young & Holsteen, 2017), according to the Web of Science Core Collection. Of these, 14 could not be accessed for classification and were excluded, leaving 1545 classifiable articles.

Among the 1545 classifiable articles, 613/1545 (39.7%) implemented a multiverse-style analysis, whereas 932/1545 (60.3%) only discussed or cited such methods without implementing them (Figure 1). The citing literature has grown steadily since 2016, comprising both articles that implemented a multiverse-style analysis and articles that only discussed or cited such methods (Figure 2 A). Among the 613 studies that implemented a multiverse-style analysis, the median number of citations was 6, and the mean 21.7 (range 0–1346; Figure 2 B); the median number of citations per year was 1.7 (mean 4.2, range 0–224.3; Figure 2 C).

**Figure 1:**
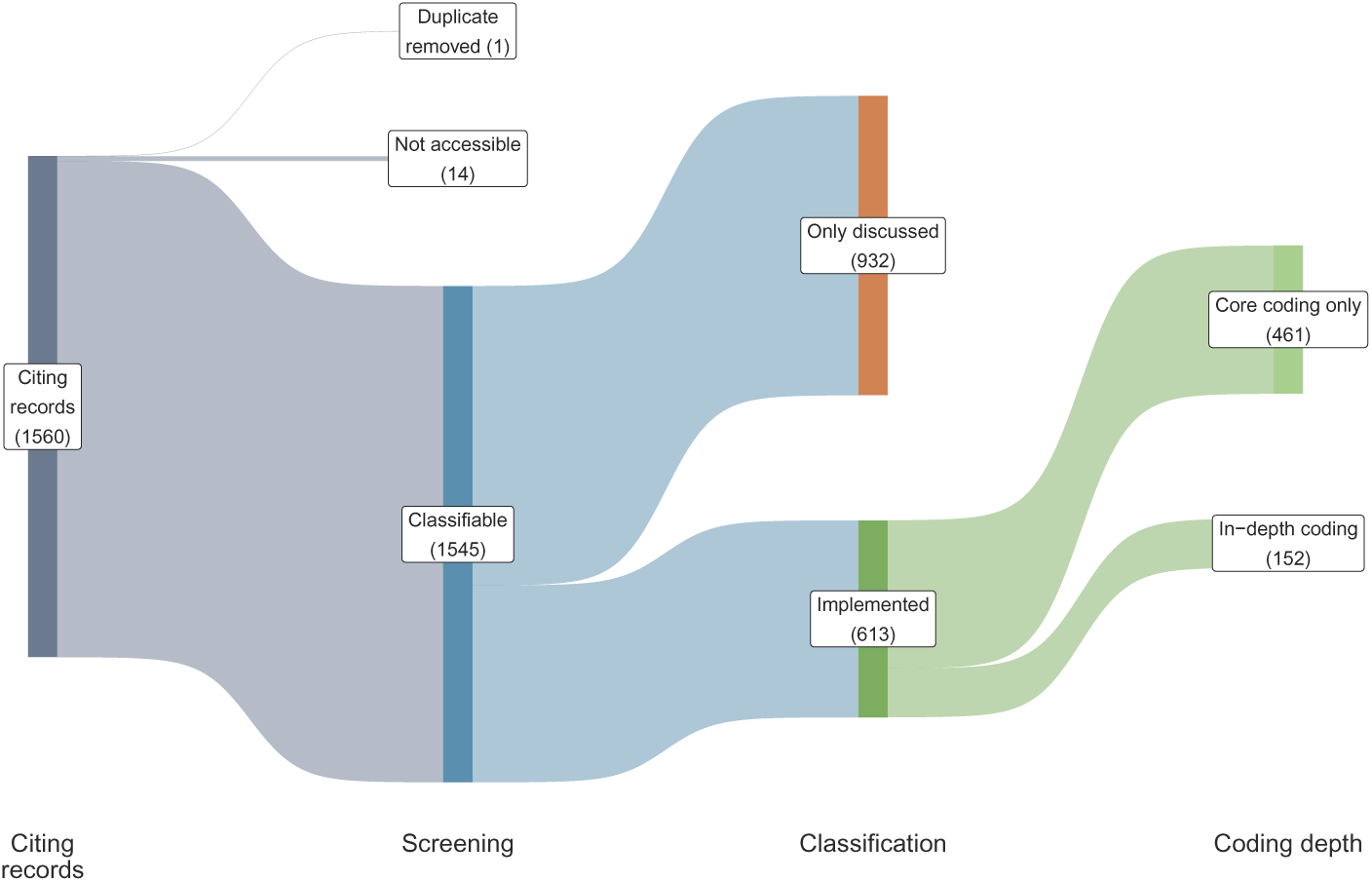
Flow of records through the assessment. Of the 1560 screened records, one was a duplicate not removed previously and 14 could not be accessed, leaving 1545 classifiable unique articles; 613 (39.7%) implemented a multiverse-style analysis and 932 only discussed or cited these methods. All 613 implemented studies were coded for the number of specifications, the decision nodes varied, and the visualization used, and a randomly selected subsample of 152 was additionally coded in depth.

**Figure 2:**
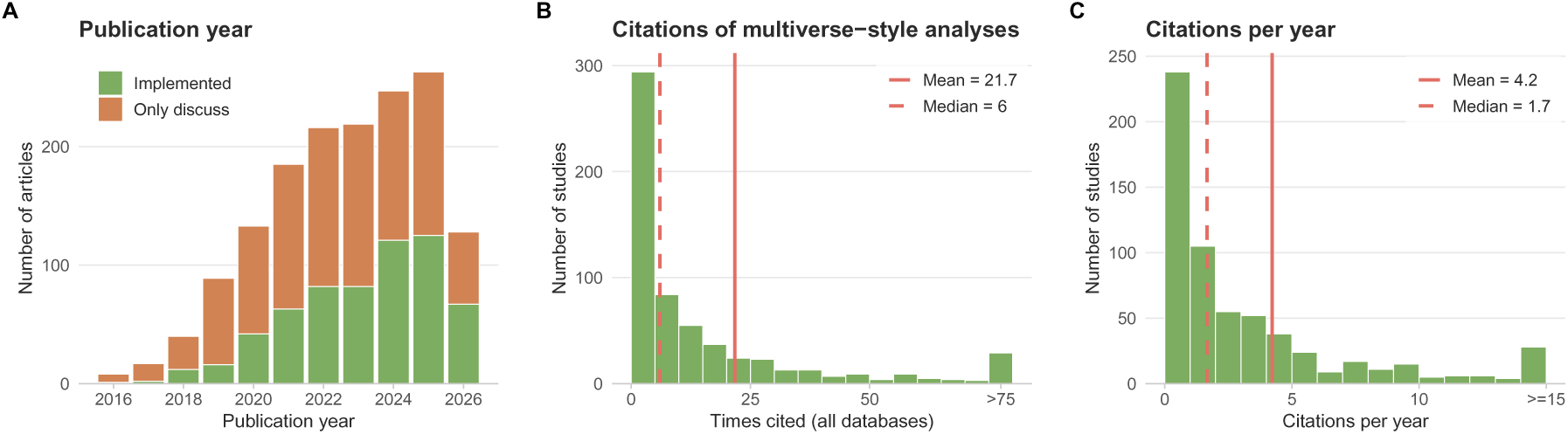
Publication years and citation impact. (A) Publication year of the 1545 classifiable articles; 2026 is partial, reflecting the mid-2026 search date. (B) Citation counts (all WoS databases) of the 613 studies that implemented a multiverse-style analysis; articles with more than 75 citations are grouped in the rightmost bin (29 articles; full range 0-1346). (C) Citations per year of the same studies (total citations divided by years since publication); studies with 15 or more citations per year are grouped in the rightmost bin.

Multiverse-style analyses were implemented across a wide range of disciplines (Table 1), most often in psychology, followed by the social sciences (sociology, political science, communication, education, and related areas) and medicine^1^. Across the 613 implemented studies, the multiverse framework (Steegen et al., 2016) was the most common (336/613, 54.8%), followed by specification curve analysis (175, 28.5%), multimodel analysis (59, 9.6%), multi-analyst (many analysts) studies (23, 3.8%), and vibration of effects (20, 3.3%). The multiverse was the most common approach in every field except economics, where specification curve analysis predominated.

**Table 1:** Fields of application of the 613 studies that implemented a multiverse-style analysis. Fields are derived from Web of Science Research Areas; a study can fall in more than one field, so percentages sum to more than 100%. ‘Framework distribution’ gives the share of each of the five frameworks (in a fixed order) among the studies in that field that had a coded framework. Framework abbreviations are MV = multiverse, SCA = specification curve analysis, MM = multimodel, MA = multi-analyst, and VoE = vibration of effects.

| Field | n | % | Framework distribution |
| --- | --- | --- | --- |
| Psychology | 244 | 39.8 | MV 76%; SCA 21%; MM 2%; MA 1%; VoE 0% |
| Social sciences | 127 | 20.7 | MV 39%; SCA 31%; MM 29%; MA 0%; VoE 1% |
| Medicine | 118 | 19.2 | MV 55%; SCA 25%; MM 4%; MA 6%; VoE 10% |
| Economics | 86 | 14.0 | MV 13%; SCA 62%; MM 18%; MA 6%; VoE 1% |
| Other/Multidisciplinary | 60 | 9.8 | MV 45%; SCA 32%; MM 3%; MA 13%; VoE 7% |
| Neuroscience | 58 | 9.5 | MV 72%; SCA 22%; MM 0%; MA 2%; VoE 3% |
| Biology | 44 | 7.2 | MV 57%; SCA 27%; MM 11%; MA 0%; VoE 5% |
| Computer science | 18 | 2.9 | MV 82%; SCA 12%; MM 0%; MA 0%; VoE 6% |
| Physical sciences & engineering | 14 | 2.3 | MV 57%; SCA 21%; MM 7%; MA 14%; VoE 0% |
| Mathematics/Statistics | 11 | 1.8 | MV 64%; SCA 18%; MM 0%; MA 9%; VoE 9% |
| Total | 613 | 100.0 | MV 55%; SCA 29%; MM 10%; MA 4%; VoE 3% |

Although for each study we coded the primary framework they cited using, many studies implementing multiverses cited several of the foundational papers together: 21.8% cited more than one, most commonly the multiverse and specification-curve papers (See Supplementary Figure 1 for a Venn diagram).

### Implementation of multiverse-style analyses

Across the implemented multiverse-style analyses, we examined how many specifications each analysis included (n = 529 studies with an extractable count) and which decision nodes were varied across those specifications (n = 613 studies). The number of specifications spanned several orders of magnitude: 88 of the 529 studies (16.6%) included 10 or fewer specifications, whereas 60 (11.3%) included 10,000 or more (Figure 3 A). For 84 studies, an explicit number of specifications could not be extracted, most often because the authors described the specification space only qualitatively (e.g., as covering all combinations of the analytical options without specifying the different options in each node). Modeling and data-processing choices were the decision nodes most frequently varied, measurement choices were considered in almost half the studies, whereas estimation choices were uncommonly varied (Figure 3 B).

**Figure 3:**
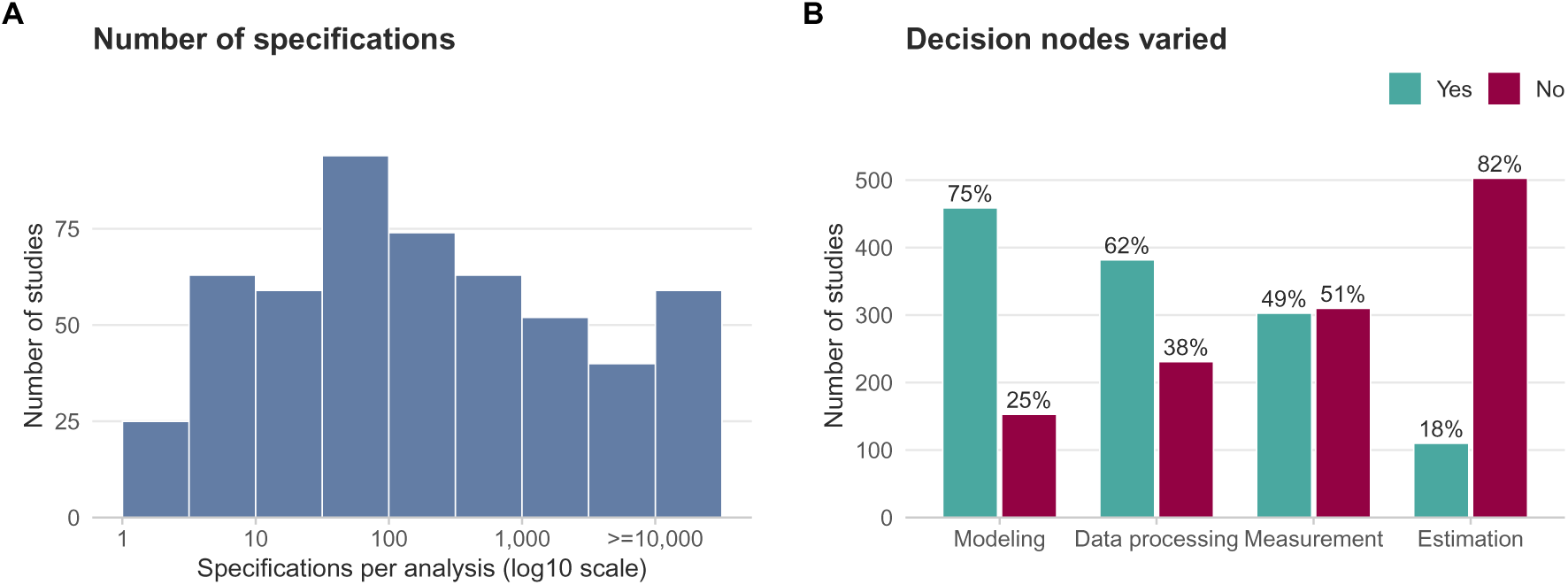
Implementation characteristics of the multiverse-style analyses. (A) Number of specifications per analysis, on a log scale (n = 529); the rightmost bar groups analyses with more than 10,000 specifications. (B) For each type of decision node, the number of studies that did (Yes) or did not (No) vary it (n = 613).

The number of specifications and the decision nodes that were varied differed both across fields of application (Table 2) and across the multiverse-style frameworks used (Table 3). By field, the median number of specifications was highest in the social sciences and economics and lowest in psychology and the physical sciences/mathematics, while modeling was the node most frequently varied in almost every field. Across frameworks the contrasts were sharper: vibration-of-effects and multimodel analyses explored by far the largest specification spaces (medians in the hundreds to tens of thousands), whereas multiverse typically varied only tens of specifications. The nodes that were varied also tracked each framework’s origins, with multimodel analyses almost exclusively varying modeling choices, specification-curve and multiverse analyses most often varying modeling and data processing, and many-analyst studies being the most likely to also vary measurement and estimation.

**Table 2:**
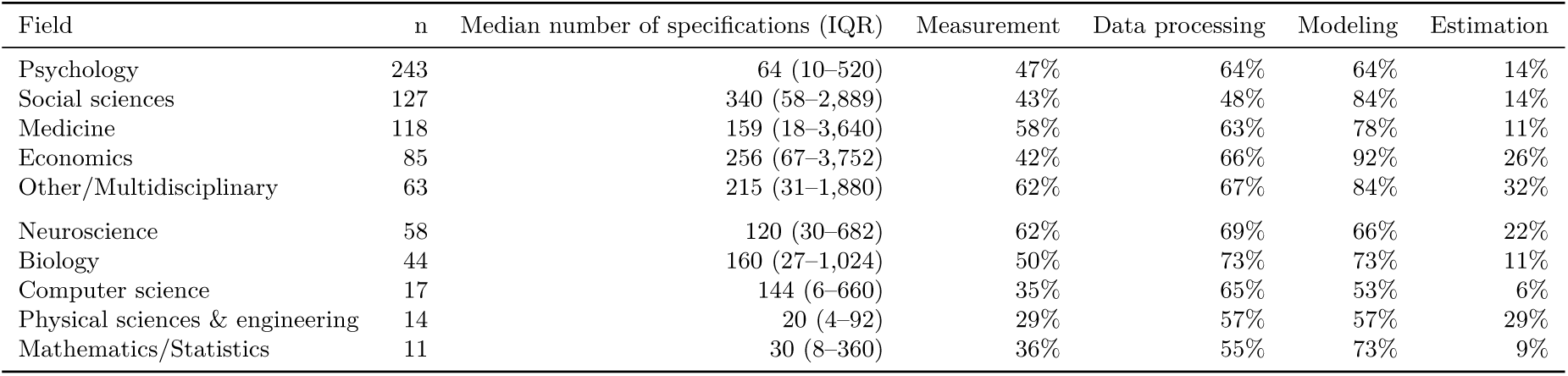
Number of specifications and decision nodes varied, by field of application. A study can be included in more than one field. Values are the median number of specifications (interquartile range) and the percentage of studies varying each decision node.

| Field | n | Median number of specifications (IQR) | Measurement | Data processing | Modeling | Estimation |
| --- | --- | --- | --- | --- | --- | --- |
| Psychology | 243 | 64 (10–520) | 47% | 64% | 64% | 14% |
| Social sciences | 127 | 340 (58–2,889) | 43% | 48% | 84% | 14% |
| Medicine | 118 | 159 (18–3,640) | 58% | 63% | 78% | 11% |
| Economics | 85 | 256 (67–3,752) | 42% | 66% | 92% | 26% |
| Other/Multidisciplinary | 63 | 215 (31–1,880) | 62% | 67% | 84% | 32% |
| Neuroscience | 58 | 120 (30–682) | 62% | 69% | 66% | 22% |
| Biology | 44 | 160 (27–1,024) | 50% | 73% | 73% | 11% |
| Computer science | 17 | 144 (6–660) | 35% | 65% | 53% | 6% |
| Physical sciences & engineering | 14 | 20 (4–92) | 29% | 57% | 57% | 29% |
| Mathematics/Statistics | 11 | 30 (8–360) | 36% | 55% | 73% | 9% |

**Table 3:** Implementation characteristics by multiverse-style framework. Columns are the five main frameworks. Rows give the number of studies, the median number of specifications (interquartile range), the percentage of studies that varied each decision node, and, among the studies in each framework that included at least one visualization, the percentage that used each visualization type.

| Characteristic | Multiverse | Specification curve | Multimodel | Multi-analyst | Vibration of effects |
| --- | --- | --- | --- | --- | --- |
| Studies (n) | 336 | 175 | 59 | 23 | 20 |
| Median specifications (IQR) | 64 (10–576) | 335 (87–1,995) | 896 (88–8,192) | 57 (15–120) | 44,980 (844–1,401,460) |
| Measurement varied | 49% | 57% | 14% | 78% | 60% |
| Data processing varied | 70% | 65% | 10% | 70% | 55% |
| Modeling varied | 63% | 89% | 98% | 91% | 70% |
| Estimation varied | 15% | 21% | 14% | 52% | 10% |
| Included at least one visualization | 62% | 94% | 56% | 87% | 85% |
| **Visualization type (% of visualizing studies)** |  |  |  |  |  |
| Specification curve | 32% | 95% | 0% | 20% | 0% |
| Forest plots/Plotting effect sizes per specification | 28% | 2% | 9% | 45% | 6% |
| Histogram/density plot | 20% | 2% | 73% | 5% | 12% |
| Outcome matrix/heatmap | 13% | 1% | 0% | 10% | 0% |
| Only tables | 8% | 1% | 18% | 0% | 0% |
| Vibration-of-effects plot | 1% | 0% | 0% | 0% | 88% |
| Boxplots | 2% | 0% | 0% | 5% | 0% |
| Multi-analysts specification curve | 0% | 0% | 0% | 20% | 0% |
| Agreement matrix (Krippendorff's ) | 0% | 0% | 0% | 0% | 0% |
| Anderson-darling statistic plot | 0% | 0% | 0% | 5% | 0% |
| Effect size per approach | 0% | 0% | 0% | 0% | 0% |
| Glasshouse plot | 0% | 0% | 0% | 0% | 0% |
| P-value threshold | 0% | 0% | 0% | 0% | 0% |
| Partial dependence plot | 0% | 0% | 0% | 0% | 0% |
| Sankey diagram | 0% | 0% | 0% | 5% | 0% |

Most implemented analyses were also accompanied by a graphical summary of the specifications: 444 of the 613 studies (72.4%) included at least one visualization. The three most common were specification curve (227 studies, 51.1%), forest plots/plotting effect sizes per specification (73 studies, 16.4%), and histogram/density plot (72 studies, 16.2%); the full distribution of visualization types is given in Supplementary Table 2. The types of visualization varied modestly across fields but the specification curve was always the most commonly used (ranging from 33% in computer science to 73% in economics). As expected, the types of visualization varied markedly depending on which methodological variant was deployed (Table 3). Most visualizing studies relied on a single type, but 20 (4.5%) combined two or more, most often histogram/density plot + outcome matrix/heatmap (4 studies).

Using a smaller sample of 152 (i.e., one-quarter) implemented analyses that we coded in greater depth, we examined several further features. The multiverse-style analysis was the study’s main analysis in 59/152 studies (38.8%) and played a secondary or supporting role in 93/152 (61.2%). The multiverse itself was preregistered in only 13/152 studies (8.6%). Few studies discussed whether their competing specifications were defensible or principled (distinguishing between equivalent, non-equivalent, or uncertain specifications) in the sense of Del Giudice & Gangestad (2021): only 6/152 (3.9%) discussed this distinction, and 3 of them judged their decisions to be of uncertain equivalence (type U decisions).

Most analyses crossed the decision nodes factorially rather than varying them one at a time: of the 152 assessed studies, 133 (87.5%) used a fully crossed (nested) design, whereas 19 (12.5%) did not; 14 because they varied only a single decision within one node and 5 because they were many-analyst studies, in which each analyst contributes an independent pipeline rather than a crossed grid. 10/152 studies (6.6%) explicitly labelled their analysis as a “limited” or “mini” multiverse.

Interpretation of the specifications was predominantly descriptive: 125/152 studies (82.2%) relied solely on a descriptive reading of the results, of which 33 (26.4%) performed explicit vote counting (counting the proportion of statistically significant specifications). A further 16/152 (10.5%) used permutation-based inference tests, and 11/152 (7.2%) applied other quantitative methods, namely robustness ratios (n = 4), meta-analysis (n = 2), analysis agreement via Krippendorff’s alpha (n = 2), and effect-variability metrics such as the relative vibration of effects, measurement vibration, or a Janus confusion matrix (n = 3).

## Discussion

In this meta-research evaluation of the literature citing six foundational papers, we examined how widely multiverse-style analyses have been adopted and how they are implemented in practice. Uptake has grown steadily since 2016, with 613 of the 1545 classifiable articles (39.7%) implementing a multiverse-style analysis. These methods have spread beyond the disciplines where they originated, although their most common fields of application remained psychology, the social sciences, and medicine. The multiverse (Steegen et al., 2016) and specification-curve (Simonsohn et al., 2020) frameworks accounted for the great majority of implementations, whereas the adoption of multi-analyst approaches remained limited, probably because they are more costly and organizationally demanding than methods where all the work and analyses can be done by a single team.

Implementation was heterogeneous. The number of specifications ranged from fewer than ten to well over ten thousand, and the decisions that were varied differed markedly across studies; modeling and data-processing choices most often, and estimation choices only rarely, although some differences could be seen across different frameworks and fields. The majority of analyses relied on a descriptive reading or vote counting of the specification set; few applied formal inference methods; and very few explicitly engaged with the defensibility or equivalence of their specifications. Preregistration of the multiverse was not common, and in most cases the multiverse analyses played only a supporting rather than a primary role.

Implementation of a multiverse analysis does not imply that the specification set was constructed in a systematic or defensible manner: a multiverse built from a narrow, non-independent, or self-confirming set of decision branches can create an illusion of robustness without meaningfully testing a finding (Del Giudice & Gangestad, 2021). Moreover, the biased construction of a technically defensible multiverse, or the selective reporting of the specifications that confirm a preferred conclusion, can subvert the very purpose these methods were designed to serve. Indeed, it can be also argued that robustness across specifications that differ in how well they are justified does not, by itself, warrant confidence in a conclusion (see Lakens et al., 2026 for a discussion of the epistemic limits of multiverse methods).

Authors of multiverse analyses may engage more explicitly in justifying the specifications included in their multiverses, adopting transparent tools such as the Systematic Multiverse Analysis Registration Tool (SMART; Short et al., 2025). In many cases, however, it may be difficult or even impossible to meaningfully justify some specifications and discard others. Running all possible specifications may then be fully appropriate or even preferable, so as to see the full range of results and examine which factors and analytical choices are most influential in shaping the conclusions. In fact, the different frameworks examined here have gone to different lengths in focusing on justified specifications versus all-inclusive analyses. The vibration of effects approach (Patel et al., 2015) typically explores all possible specifications, whereas specification-curve (Simonsohn et al., 2020) studies usually try to justify their selective choices.

Methodologists can develop concrete guidance and tools for principled, non-redundant specification-space construction, lowering the practical cost of doing this well. Finally, future studies could assess the defensibility and equivalence of the pipelines used in the studies reviewed here. Moreover, when specific results are chosen as more likely to represent the data, their replicability should be assessed against subsequent replication studies in independent samples and new data.

Our study has limitations inherent to its design. Because we identified studies through citations to specific, highly cited foundational papers, we necessarily missed analyses that implement equivalent ideas without citing these specific papers. Each of these methods has been presented in additional methodological papers which have also been cited, albeit less frequently than those we used as indices for our searches. Some of these methodological variants have also been presented in earlier work under different terminology (e.g., the vibration of effects concept was introduced as a “vibration ratio” in a 2008 paper (Ioannidis, 2008), seven years before the index paper that we use here) or were inspired by earlier work (e.g., the highly influential and extremely highly cited paper “I Just Ran Two Million Regressions” by Xavier Sala-i-Martin, published in the American Economic Review in 1997 (Sala-i-Martin, 1997)). Therefore, our figures clearly underestimate the total uptake of these methods across the various domains of the scientific literature, but they define a reproducible cohort of relevant studies. It is nevertheless not known whether studies using such ideas without citing any of the six papers that we selected may also have different features than those of the studies assessed here. Another limitation is that in-depth coding of some implementation features was carried out on a subsample by a single coder, so some estimates may carry measurement error. However, the range of frequencies for the features assessed was still highly revealing of the lack of some standard research practices.

Eventually, different variants of multiverse-style analyses are likely to become increasingly used. Documenting their growing prevalence and how exactly they are deployed may help accelerate their careful and nuanced adoption. In the current evaluation of a large number of relevant papers, we have highlighted practices that authors may consider to ensure that these analyses genuinely achieve their intended purpose. We also hope this coding scheme will be reapplied rather than treated as a one-time snapshot, so that the field’s uptake of principled robustness assessment can be tracked as future developments take place.

## Acknowledgements

This evaluation was not preregistered and it is exploratory of the field. During the preparation of this manuscript, the authors used Claude Opus 4.8 for code development and manuscript editing.

## Data and Code Availability

All data, the analysis scripts required to reproduce the analyses, and the source files needed to generate this fully reproducible manuscript are openly available on the Open Science Frame-work (https://osf.io/j7nsf/).

## Supplementary Materials

### Classification of Research Areas into fields

**Supplementary Table 1.** Mapping of Web of Science Research Areas to the field categories used in this review. An article was assigned to a field when any of its Research Areas contained one of the listed terms (case-insensitive substring match); the categories were evaluated in the order shown, and articles matching none of the terms were assigned to Other/Multidisciplinary.

| Field | Web of Science Research Areas (matched terms) |
| --- | --- |
| Neuroscience | Neurosci, Neurology |
| Psychology | Psychology, Behavioral sciences |
| Computer science | Computer science, Informatics, Information science, Robotics, Automation & control, Telecommunications, Imaging science |
| Economics | Business & economics, Economics, Operations research |
| Social sciences | Sociology, Social sciences, Social work, Communication, Criminology, Demography, Family studies, Anthropology, Urban studies, Ethnic studies, Women, Development studies, Social issues, Public administration, Cultural, Government & law, International relations, Education, Area studies, Biomedical social |
| Medicine | Medicine, Medical, Health, Psychiatry, Public, environmental, Radiology, Oncology, Geriatric, Nutrition, Pharmacolog, Cardiov, Surgery, Anesthes, Ophthalm, Rehabilitation, Endocrin, Infectious, Pediatric, Orthopedic, Obstetric, Dentistry, Respiratory, Urology, Nursing, Immunology, Emergency, Pathology, Rheumatology, Otorhino, Substance abuse, Audiology, Sport sciences, Life sciences & biomedicine |
| Mathematics/Statistics | Mathematic, Statistics |
| Biology | Ecology, Environmental sciences, Biology, Biochem, Physiology, Agriculture, Genetics, Zoology, Marine, Biodiversity, Microbiology, Reproductive, Anatomy, Veterinary, Food science, Plant, Biotechnology, Cell biology, Fisheries, Forestry, Entomology |
| Physical sciences & engineering | Physics, Chemistry, Geolog, Meteorolog, Atmospheric, Water resources, Oceanograph, Optics, Acoustics, Astronomy, Geochem, Mineral, Engineering, Materials science, Energy & fuels, Transportation, Construction, Instruments, Mechanics, Remote sensing |
| Other/Multidisciplinary | Fallback: articles whose Research Areas matched none of the terms above |

### Visualization types

**Supplementary Table 2.**
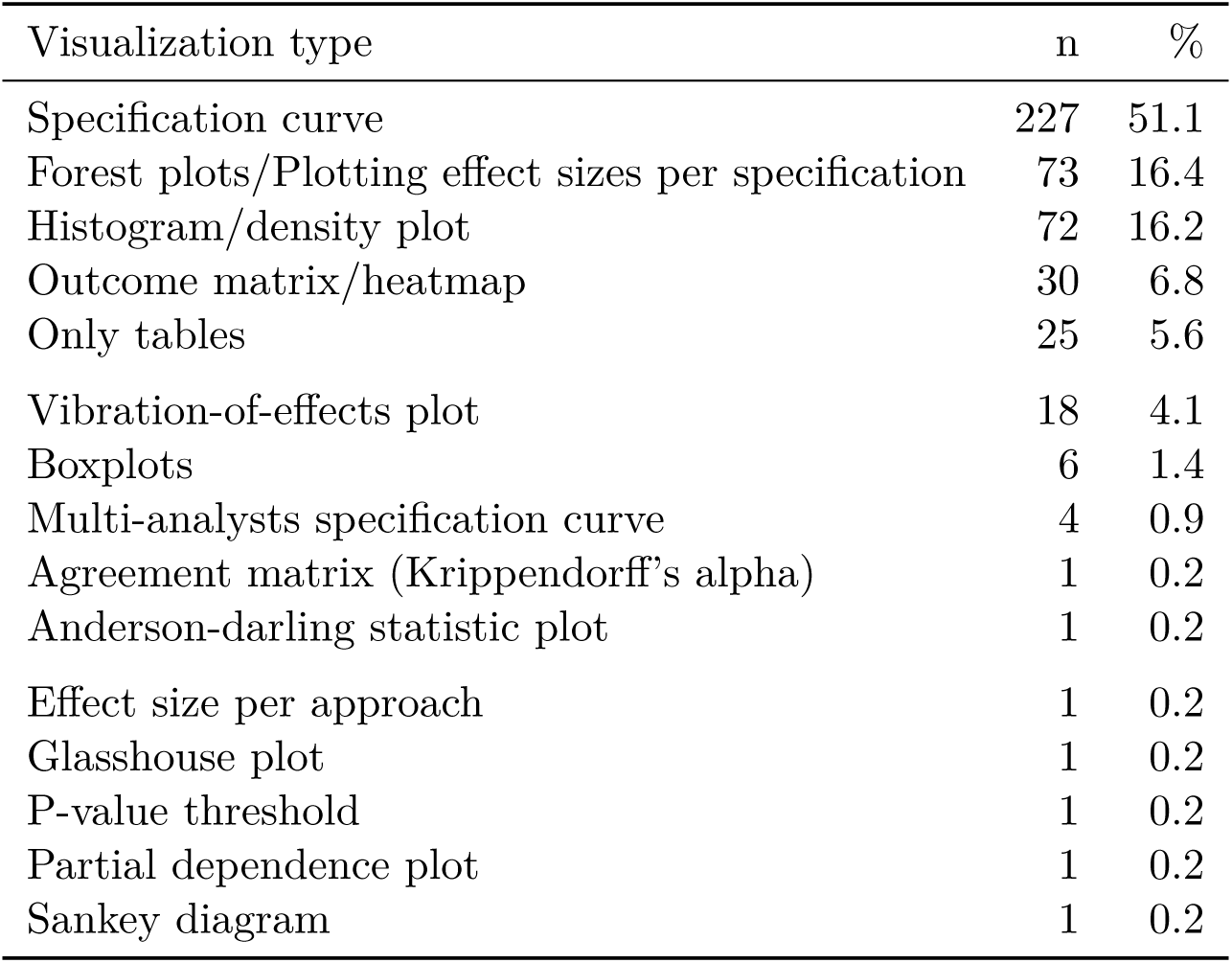
Visualization types used by the 444 studies that included at least one visualization. Because some studies used more than one type, counts sum to more than 444 and percentages (of the 444 visualizing studies) to more than 100%.

### Co-citation of the foundational papers

**Supplementary Figure 1.**
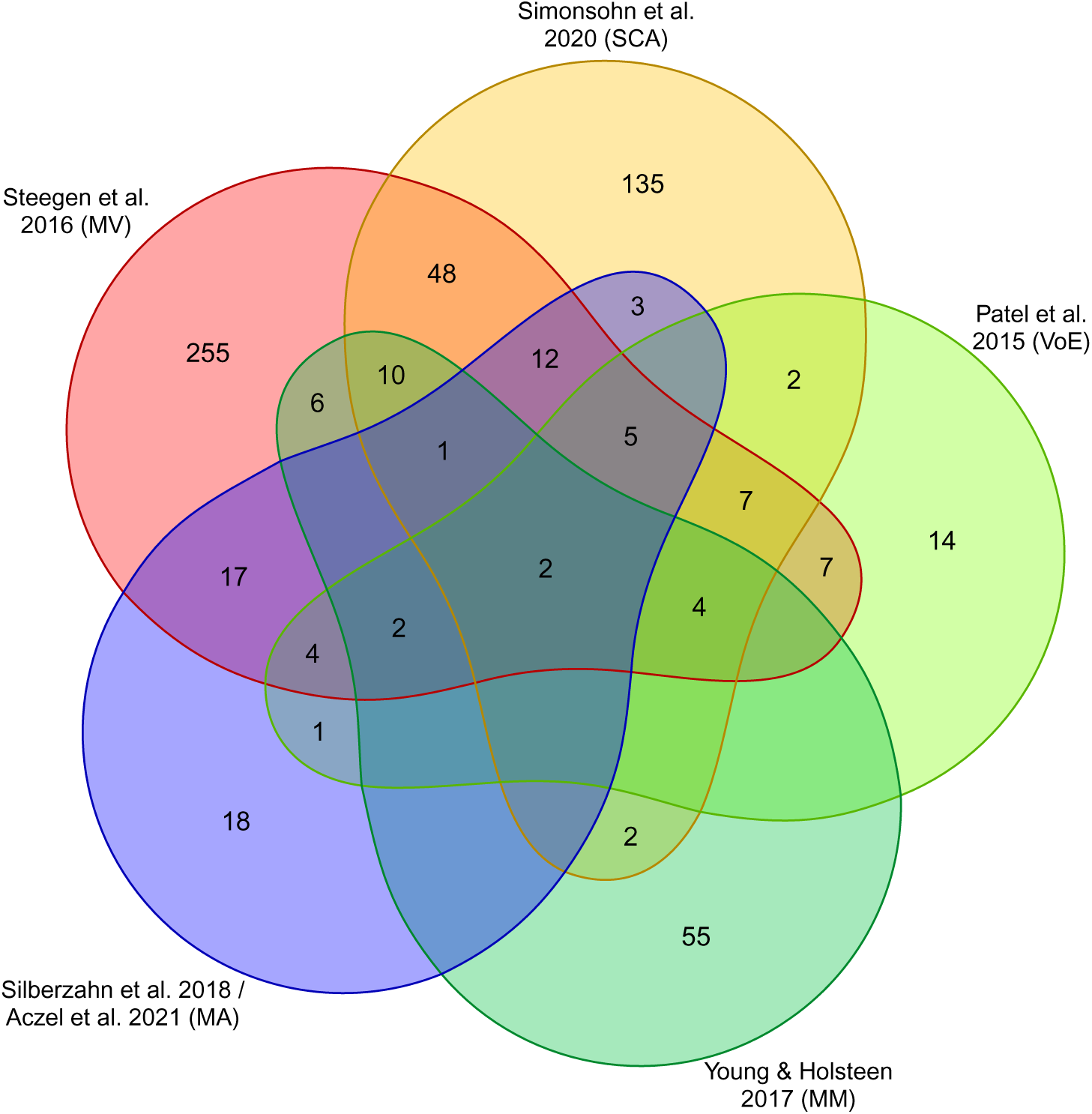
Co-citation of the six foundational papers among the 610 implemented studies whose DOI could be matched to the Web of Science citation records (3 of the 613 implemented studies had no DOI in the citation export and could not be matched). This analysis is based on the citation records: each study is assigned to a set according to which of the six foundational papers it *cited* (not by the framework it was coded as using), and overlaps are studies that cited several. Each set is labelled by the foundational reference and, in parentheses, the corresponding framework abbreviation (MV = multiverse, SCA = specification curve analysis, VoE = vibration of effects, MM = multimodel, MA = multi-analysts); the MA set combines the two multi-analyst foundational papers (Silberzahn et al. 2018 and Aczel et al. 2021).

## Footnotes

1 More information about the classification of each field can be found in Supplementary Table 1.

